# HOCl Forms Lipid *N*-Chloramines in Cell Membranes of Bacteria and Immune Cells

**DOI:** 10.1101/2024.06.25.600395

**Authors:** Lisa R. Knoke, Sara Abad Herrera, Sascha Heinrich, Natalie Lupilov, Julia E. Bandow, Thomas Günther Pomorski

## Abstract

Neutrophils orchestrate a coordinated attack on bacteria, combining phagocytosis with a potent cocktail of oxidants, including the highly toxic hypochlorous acid (HOCl), renowned for its deleterious effects on proteins. Here, we examined the occurrence of lipid *N*-chloramines *in vivo*, their biological activity and neutralization. Using a chemical probe for *N*-chloramines, we demonstrate their formation in the membranes of bacteria and monocytic cells exposed to physiologically relevant concentrations of HOCl. *N*-chlorinated model membranes composed of phosphatidylethanolamine, the major membrane lipid in *Escherichia coli* and an important component of eukaryotic membranes, exhibited oxidative activity towards the redox-sensitive protein roGFP2, suggesting a role for lipid *N*-chloramines in protein oxidation. Conversely, the cellular antioxidant glutathione neutralized lipid *N*-chloramines by removing the chlorine moiety. We propose that lipid *N*-chloramines, like protein *N*-chloramines, are involved in inflammation and accelerate the host immune response.

## Introduction

Professional phagocytotic cells, such as neutrophils, are among the first lines of defense in the mammalian immune system. When pathogens encounter these cells, they are phagocytized and killed in the so-called phagolysosome. In this compartment, pathogens are exposed to a complex mixture of lytic proteins, toxins and oxidative and nitrosative species (Furtmüller et al., 1998; Ulfig and Leichert, 2021). The release of reactive oxygen (ROS) and nitrogen (RNS) species is referred to as oxidative burst and is initiated by the NADPH oxidase NOX2 at the phagosomal membrane, which transfers electrons onto molecular oxygen resulting in the formation of superoxide radicals. Within the phagolysosome, this superoxide radical is further converted to hydrogen peroxide (H_2_O_2_). Using H_2_O_2_ and Cl^-^ anions, the myeloperoxidase (MPO) catalyzes the production of the highly toxic oxidant hypochlorous acid (HOCl) (Sultana et al., 2020; Winterbourn, 2008). It has been shown that upon phagocytosis, bacterial proteins containing free thiol are oxidized within seconds (Degrossoli et al., 2018; Xie et al., 2020, 2019). In addition to thiol oxidation, HOCl attacks amino groups in various target molecules such as lysine residues in proteins, aminosugars and nucleic acids, resulting in *N*-chlorination (Carr et al., 2001; Hawkins and Davies, 2002; Panasenko et al., 2013; Rees et al., 2005; Varatnitskaya et al., 2022; Winter et al., 2008).

Hypochlorous acid is a very weak acid and that exists as a mixture of ^-^OCl and HOCl (pKa = 7.59) (Morris, 1966) at physiological pH. It is suggested that only the uncharged, protonated form can cross cell membranes (Kawai et al., 2006; Zavodnik et al., 2001). When phagocytized bacteria are exposed to HOCl, their membrane is one of the first targets. In many bacterial organisms, phosphatidylethanolamine (PE) is a major membrane lipid, containing a diverse mixture of saturated and unsaturated fatty acids, and is also relevant in eukaryotes. The membranes of the model organism *Escherichia coli* consist of 70-75% PE, 10-15 % phosphatidylglycerol (PG) and 5-10% cardiolipin (CL) (Kleetz et al., 2021). In a study by Carr et al. (1998) using liposomes prepared from *E. coli* phospholipids, HOCl modified PE either by *N*-chlorination of the head group amine or by chlorohydrin formation at the C=C double bond in unsaturated fatty acids. Predominantly, PE *N*-chloramines were formed until all free amine groups were consumed, after which chlorohydrins began to accumulate (Carr et al., 1998).

Recent studies with proteins revealed that HOCl induced *N*-chlorination of basic lysine and arginine amine groups mediates the activation of a holdase-like chaperone function in several bacterial proteins such as RidA (Müller et al., 2014; Varatnitskaya et al., 2022) or CnoX (Goemans et al., 2018) and human proteins such as serum albumin (HSA) (Ulfig et al., 2019). Furthermore, *N*-chlorinated HSA activates ROS production in immune cells and antigen presentation (Ulfig et al., 2021, 2019). In contrast to proteins, most of our knowledge about lipid *N*-chloramines is based on *in vitro* studies. Even more recent reviews on HOCl targets (Andrés et al., 2022; Snell et al., 2022) do not provide information on lipid *N*-chloramines *in vivo*. A plausible reason for the lack of available data is that *N*-chloramines are relatively unstable due to their high reactivity (Fuchs et al., 2011; Pattison et al., 2003). Although detection by thin layer chromatography (TLC) (Carr et al., 1998; Richter et al., 2008) or detection of oxidation products by MS (Flemmig et al., 2009) is possible, reliable detection methods for lipid *N*-chloramines *in vivo* are still lacking (Fuchs, 2014). A few studies even report direct detection of PE *N*-chloramines *in vitro* by ESI-MS (Guo et al., 2021; Kawai et al., 2006) or MALDI-TOF MS (Jaskolla et al., 2009). Therefore, a reliable detection method for lipid *N*-chlorination is needed in order to understand the physiological relevance of this modification.

## Materials and methods

### Strains, plasmids and growth conditions

The strains and plasmid used in this study are listed in Table 1. *E. coli* BW25113 carrying pCC_*roGFP2* plasmid was grown at 37 °C in Luria-Bertani (LB) medium, supplemented with antibiotics for plasmid maintenance (200 µg/mL ampicillin), unless otherwise specified.

**Table 1.**
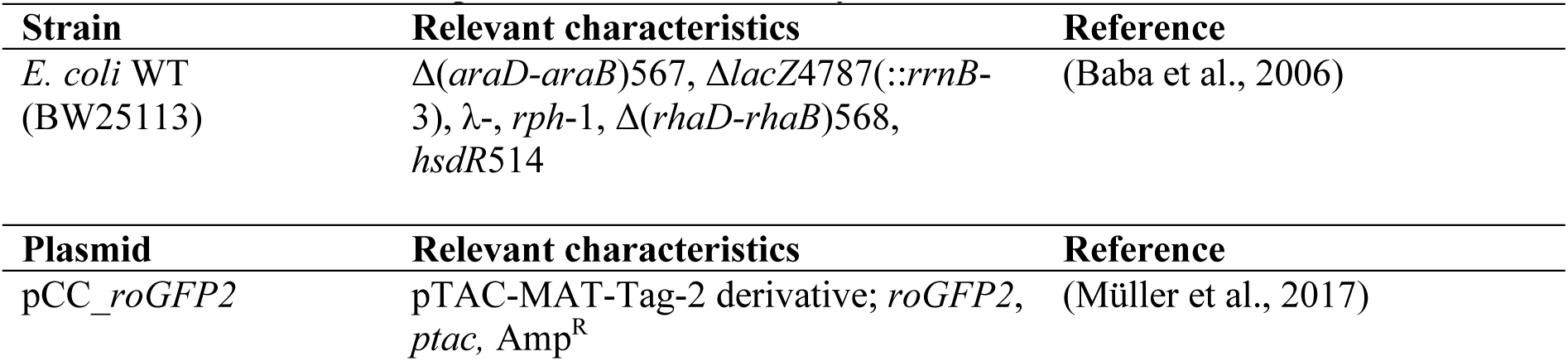
Bacterial strain and plasmid used in this study.

Expression of roGFP2 was induced with 0.2 mM IPTG (isopropyl β-D-thiogalactopyranoside) at an OD_600_ of ∼0.6–0.8, followed by 16 h incubation at 20 °C. The cells were then used for HOCl treatment and analysis of lipid *N*-chlorination. THP1 cells were cultured in RPMI medium supplemented with phenol red (#61870036 ThermoFischerScientific, Darmstadt Germany) and 10 % FCS (fetal calf serum, Brazil) #10270106, ThermoFischerScientific, Darmstadt Germany) at 37 °C, 5 % CO_2_ and 95% relative humidity. After 3-4 days of cultivation, the medium was completely changed and the cell number was adjusted to 200,000 cells/mL.

### Preparation of dansyl reagents for labelling

The derivatizing agents DANSCl (dansyl chloride) and DANSO_2_H (dansyl sulfinic acid) were prepared as previously described with minor changes (Varatnitskaya et al., 2022). A 200 mM DANSO_2_H solution was freshly prepared by dissolving the powder in 200 mM NaHCO_3_ and adjusting the pH to 9 with 1 M NaOH. The 200 mM DANSCl (abcr GmBH, Karlsruhe, Germany) stock solution was prepared by dissolving the powder in acetone.

### Preparation of <u>L</u>arge <u>U</u>nilamellar <u>V</u>esicles (LUV)

For the preparation of liposomes, a thin lipid film was generated, rehydrated followed by freeze-thawing and manual extrusion. All lipids were purchased from Avanti Pola Lipids (Alabaster, AL). The fully saturated lipid DMPC (14:0 PC; 1,2-dimyristoyl-*sn*-glycero-3-phosphocholine) or a DMPC/DMPE (14:0 PE; 1,2-dimyristoyl-*sn*-glycero-3-phosphoethanolamine) lipid mixture (30/70 molar ratio) was dissolved in chloroform/methanol (9/1, v/v) and mixed in a Pyrex/Duran glass vial with a total lipid amount of 7 µmols. Lipids were dried overnight at 250 mbar in a rotary evaporator at room temperature. The lipid films were re-hydrated in 1 mL PBS buffer (NaCl 150 mM, 1 mM KH_2_PO_4_, 3 mM Na_2_HPO_4_-7xH_2_O, pH 7.4) preheated to 65 °C (phase transition temperature of DMPE and DMPC, 50 and 24 °C), facilitated by vigorous mixing in the presence of a glass bead (5 mm diameter) for 10 min, resulting in a final lipid concentration of 5-7 mM. Vesicles composed of 100 % PC are referred to as PC vesicles and vesicles composed of 70 mol% PE and 30 mol% PC are referred to as PE vesicles. The vesicles were subjected to ten freeze/thawing cycles in liquid nitrogen (1 min) and 65 °C (2 min). Liposomes were extruded over a stack of two polycarbonate membranes (pore size 400 nm) 21 x times at 65 °C using a mini-extruder (Avanti Polar Lipids) according to the manufacturer’s instructions. Lipid concentration after extrusion was determined by phosphate quantification. The extruded liposomes were stored in a glass vial at 4°C for a maximum of four days.

### Determination of lipid phosphorous

The phospholipid phosphorus was quantified after heat destruction in the presence of perchloric acid by standard techniques as described previously (Chifflet et al., 1988). Briefly, 650 µL perchloric acid (<72 %, v/v) was added to the liposome solution (5 to 20 µL); For controls, PBS or NaHCO_3_ were used. Samples were heated at 195 °C until solution appeared clear (approximately 2-3 hours), then cooled to room temperature and adjusted to 250 µL with ddH_2_O. Subsequently, 3.3 mL ddH_2_O, 0.5 mL freshly prepared ammonium molybdate solution (2.5 %, w/v) and 0.5 mL ascorbic acid (10 %, w/w) were added. Inorganic phosphate (20 to 100 nmol in 250 µL ddH_2_O) were used as standard. After thorough mixing and heating at 80 °C followed by cooling on ice, absorbance at 812 nm was measured in a multiplate reader (CLARIOStar Plus, BMG LABTECH GmBH, Germany).

### Chlorination and dansyl labelling of phosphatidylethanolamine in model membranes

Vesicle chlorination followed the protocol established for the RidA protein (Varatnitskaya et al., 2022). Vesicle solutions were adjusted to 3 mM total lipid concentration. HOCl treatment was conducted using 900 nmol total lipid (300 µL of 3 mM). HOCl was then added to both, PC and PE vesicles at equimolar molar ratio to PE (2.1 mM, 630 nmol HOCl). Water-treated vesicles served as negative control. After incubation at 37 °C for 10 min, excess HOCl was removed and the buffer replaced with 200 mM NaHCO_3_, pH 9, using a NAP-5 column (Cytiva). The lipid concentration after HOCl removal was 0.8 mM (0.56 mM PE) in 1 mL. Chlorinated vesicles were split in aliquots containing 100 nmols total lipid and incubated with a two-fold molar excess of DANSCl or DANSO_2_H (140 nmol) relative to the PE amount (70 nmol) for 10 min at 37 °C with shaking at 600 rpm. Water-treated vesicles were used as labelling specificity control and unlabelled vesicles as negative control. Finally, lipids were isolated and analyzed by TLC for dansyl derivatization (see below). Water-treated and unlabelled vesicles served as controls.

### Chlorination and dansyl labelling of phosphatidylethanolamine in E. coli and THP1 membranes

*E. coli* BW25113 (WT) cells expressing roGFP2 were harvested, washed three times in PBS, and adjusted to an OD_600_ of 6.0 in PBS. To estimate the bacterial PE content, the following assumptions were made: an OD_600_ of 1.0 corresponds to 8 x 10^8^ cells/mL, each cell weighs approximately 3 x 10^-13^ g (dry weight) and approximately 10 % of the dry weight is lipids. With the molar weight of PE of about 715 g/mol and based that 75 % of all *E. coli* lipids are PE, we estimated a PE concentration of 320 µM in a bacterial solution with an OD_600_ of 6. Aliquots of 2 mL cell suspension were treated with either ddH_2_O (no chlorination), a 1/1 molar ratio of HOCl/PE (320 µM HOCl), or a 10-fold molar excess of HOCl/PE (3.2 mM HOCl) for 10 min at 37 °C. Excess HOCl was removed by centrifugation (1 min, 20,000 x g, 4 °C) and two PBS washes. Cells were resuspended in 2 mL 200 mM NaHCO_3_ (pH 9) and divided into three aliquots, each treated with either 640 µM DANSCl, 640 µM DANSO_2_H (2/1 molar ratio with respect to PE) or ddH_2_O for 10 min at 37 °C with shaking at 600 rpm. From these solutions 100 µL were used directly for lipid analysis. For THP1 cells, we estimated a dry weight of 1.25 x 10^-9^ g per cell, with lipids comprising 10 % of this mass. Assuming 30 % of the total lipids are PE (molar mass of about 715 g/mol), we estimated a PE concentration of 0.5 mM in a 10^6^ cells/mL solution. For HOCl treatment, 800 µL of 10^6^ cells/mL solution in PBS were incubated with either 0.5 mM (1x HOCl) or 5 mM (10x HOCl) for 10 min at 37 °C. Cells were then washed twice with 1 mL PBS and resuspended in 800 µL NaHCO_3_ (pH 9). Water served as negative control. Next, 1 mM DANSO_2_H or DANSCl (2/1 molar ratio in relation to PE) were added to 260 µL cell suspension and incubated for 10 min at 37 °C before lipid isolation. Water-treated cells were used as a negative control for chlorination.

### DTT and GSH treatment of chlorinated vesicles

For chlorination, 1050 nmol HOCl (equimolar relative to PE) was added to 1500 nmol of lipids in PE vesicles (500 µL, 3 mM). Excess HOCl was removed as described above using NAP-5 columns, with a recovery of 90 % total lipid (1350 nmol) and a lipid concentration of 1.13 mM in 1.2 mL due to inherent dilution in NAP-5 chromatography. Aliquots containing 100 nmol lipids (70 nmol PE) were labelled with a two-fold molar excess of DANSCl or DANSO_2_H (140 nmol) relative to PE (chlorinated sample), using water as a negative control. To remove chlorine from lipid head groups, a 10-fold molar excess of DTT (Dithiotreitol) or GSH (reduced glutathione) (5.17 µmol) was added to 517 nmol total lipids, and the vesicles were incubated for 10 min at 37 °C with shaking at 600 rpm. Excess DTT was removed, and buffer was exchanged to 200 mM NaHCO_3_ (pH 9) using NAP-5 columns. For GSH removal, centrifugation (10 min, 20,000 x g) followed by washing with NaHCO_3_ was used. After reductant removal, an assumed 90% recovery yielded a lipid concentration of 0.39 mM. Aliquots containing 100 nmols total lipids were derivatized with DANSCl or DANSO_2_H (140 nmol) using a two-fold molar excess relative to PE (70 nmols). Unlabelled vesicles served as negative controls. All lipids (100 nmol for each step) were isolated and analyzed by TLC.

### Lipid analysis

Lipids were isolated using a modified Bligh and Dyer protocol (Bligh and Dyer, 1959; Knoke et al., 2020). To 100 µL of liposome solution, 375 µL of a chloroform/methanol (1/2, v/v) were added, mixed and incubated for 5 min at room temperature. Next, 100 µL chloroform and 100 µL ddH_2_O were added, mixed and samples were centrifuged for 5 min at 20,000 x g to improve phase separation. For larger volumes, organic solvent volumes were adjusted proportionally. The chloroform phase was collected, dried in a SpeedVac for 20 to 40 min and the lipid film was dissolved in 30 µL of chloroform/methanol (9/1 for DMPE samples, 1/2 for others; v/v) and applied on glass backed silica gel plates (60, 10 × 20 cm; Merck, #1.05626.0001). Plates were developed in chloroform/ethanol/triethylamine/ddH_2_O 30/35/35/7 (v/v/v/v) for 45 min, dried for 15 min, and dansylated lipids were detected via UV transillumination. Primulin (0.005 % (w/v) in 80 % acetone (v/v)) was sprayed on the plate to visualize all lipids under UV light.

### Sample preparation for MS analysis and LC-coupled MS^E^ measurements

For mass spectrometry, PE vesicles (70 % PE, 30 % PC) were chlorinated with HOCl as described above. Excess HOCl was removed, and buffer exchange was performed using NAP-5 columns. Lipids (175 nmol) were labelled with a two-fold molar excess of DANSCl or DANSO_2_H (245 nmol), relative to the PE amount (122.5 nmol), and separated via TLC. Water-treated vesicles served as control for *N*-chlorination, and unlabelled vesicles as negative control for dansyl labelling. Lipid spots showing dansyl fluorescence were scraped from the TLC plate and transferred to glass vials. To remove residual silica, 90 µL of ddH2O were added and mixed vigorously, followed by the addition of 160 µL chloroform and 320 µL methanol with mixing at each step. Samples were incubated for 5 min at room temperature, then 275 µL of ddH_2_O were added and mixed. After centrifugation at 500 x g for 5 min at room temperature, the upper phase was discarded, and the bottom phase was transferred in a fresh vial. This process was repeated once to ensure complete removal of silica. Lipids were dried overnight in an evaporator at 250 mbar and dissolved in 30 µL methanol/chloroform (1/9, v/v) for MS analysis.

Of these samples 5 µL, were injected on an ACQUITY UPLC I-Class System (Waters, Milford, Massachusetts) equipped with an ACQUITY UPLC-BEH C18 column (particle size 1.8 µm, column dimensions: 2.1 x 50 mm, Waters). A gradient with 10 mM ammonium acetate in H_2_O (A) and 10 mM ammonium acetate in 1:1 acetonitrile/isopropanol (B), each with 0.1 % acetic acid, was used with a flow rate of 0.3 ml/min (Pi et al., 2016) (Table 2).

**Table 2.**
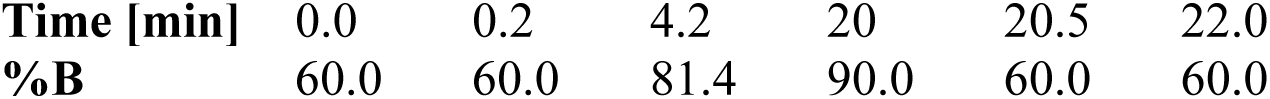
Liquid chromatography gradient.

Data-independent MS^E^ measurements were performed with a Vion IMS QToF (Waters) with an ESI source in negative sensitivity mode. Masses in a range of 50 to 1000 m/z were detected with 0.2 s per scan and leucine enkephalin being injected as a reference mass every 5 min. Used parameters: capillary voltage 0.8 kV, sample cone voltage 40 V, source offset voltage 80 V, cone gas flow 50 l/h, desolvation gas flow 1000 l/h, source temperature 120 °C, desolvation temperature 500 °C, collision gas N_2_, collision low energy 6 V, collision high energy ramp 28-63 V.

Peak picking and lock mass correction were performed using UNIFI (Waters, version 1.9.13.9) with the following parameters: automatic chromatographic peak width, automatic peak detection threshold, low energy intensity threshold 200 counts, high energy intensity threshold 500 counts, chromatographic peak width to apply during cluster creation 0.5, chromatographic peak width to apply during high to low energy association 0.5, drift peak width to apply during cluster creation 0.5, drift peak width to apply during high to low energy association 0.5, intensity threshold to apply during high to low energy association 50, maximum considered charge 1, maximum number of isotopes 5, minimum allowed monoisotopic to largest isotope ratio 0.7, allow wider tolerance for saturated data.

Method parameters for identification: retention time tolerance 0.15 min; target match tolerance 10 PPM, CCS tolerance 2 %.

### Determination of the redox state of roGFP2 in the cytosol of E. coli after HOCl exposure

The redox state of roGFP2 expressed in the cytosol of *E. coli* WT carrying pCC_*roGFP2* plasmid was determined as described previously (Degrossoli et al., 2018; Gutscher et al., 2008; Xie et al., 2020). Briefly, roGFP2 expression was induced as described above for 16 h at 20 °C. Cells were harvested, washed twice with PBS and adjusted to an OD_600_ of 6.0 in PBS. Excitation spectra (350-500 nm) with fixed emission at 510 nm (5 nm bandwidth) were recorded using an FP-8500 spectrofluorometer (Jasco, Tokyo, Japan) at 25 °C with continuous stirring for at least 1 min before adding HOCl at a one-fold (320 µM) or ten-fold (3.2 mM) molar excess relative to the theoretical PE concentration. Spectra were recorded for at least 5 min.

The fluorescence excitation ratios (405/488 nm) were used to calculate the oxidation state of the probe (Lohman and Remington, 2008). Cells treated with 1 mM AT-2 (aldrithiol-2) or 10 mM DTT (dithiotreitol) served as controls for complete oxidation or reduction of the probe. Equation [1] was used for calculation.

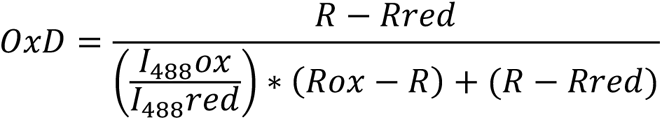

With *Rox* being the 405/488 nm ratio of oxidized and *Rred* of reduced roGFP2. *I_488_ox* and *I_488_red* represent the fluorescence intensities of roGFP2 at 488 nm under oxidizing or reducing conditions, respectively. *R* is the measured 405/488 nm ratio of roGFP2 in the respective sample.

### roGFP-2 oxidation by chlorinated vesicles

The purified redox-active roGFP2 protein was reduced as described previously (Knoke et al., 2023). The concentration was then adjusted to 0.2 µM, and the excitation spectra (350 nm to 500 nm, fixed emission at 510 nm, bandwidth: 5 nm) were recorded in an FP-8500 spectrofluorometer (Jasco, Tokyo, Japan) at 25 °C with continuous stirring for at least 1 min. Oxidized (0.1 mM AT-2) or reduced (2 mM DTT) roGFP2 served as controls for calculating the degree of oxidation. To test the oxidative activity of chlorinated vesicles, PC or PE vesicles were prepared and chlorinated with 2.1 mM HOCl for 10 min. Excess oxidant was removed using a NAP-5 column. A 100-fold molar excess of chlorinated PE (26 µM total lipids, 20 µM PE) was added to the reduced roGFP2 protein (0.2 µM) in PBS. Excitation spectra were then recorded for at least 5 min, followed by the addition of 20 mM DTT to completely reduce the sensor. Untreated PE-containing vesicles served as negative control. To exclude probe oxidation in the presence of lipids in general, PC vesicles treated with 2.1 mM HOCl or water were used. HOCl-treated PBS (2.1 mM HOCl in PBS) that passed an NAP-5 column was added to rule out residual HOCl in the chlorinated samples. Data was evaluated as described above [Equation 1].

## Results

### Formation and detection of N-chloramines in phosphatidylethanolamine upon HOCl exposure in model membranes

During phagocytosis, neutrophils produce HOCl to kill engulfed pathogens in the phagolysosome, resulting in various modifications of different biomolecules, including *N*-chloramines (Hawkins and Davies, 2002; Varatnitskaya et al., 2022). When bacteria encounter HOCl in the phagolysosome, one of the first targets are membrane phospholipids, most importantly PE (Kleetz et al., 2021). PE is prone to *N*-chlorination at its amine head group (Carr et al., 1998). It is unknown whether PE *N*-chloramines occur *in vivo* and their physiological role and biological activity are still under investigation, since, due to the inherent instability of *N*-chloramines, their detection is challenging. Recently, the dansyl derivative dansyl sulfinic acid (DANSO_2_H) was successfully used to stabilize and identify *N-*chlorinated lysine residues in the model protein RidA after HOCl exposure (Varatnitskaya et al., 2022). The reaction of DANSO_2_H with *N*-chloramines produces dansyl chloride (DANSCl) and a free amine, which further react to a stable, fluorescent sulfonamide. DANSCl is a well-established reagent that reacts with free amino groups of proteins (Chen, 1968) and lipids (Schindler and Teuber, 1978; Shechter et al., 1971). We hypothesized that DANSO_2_H would also be suitable for the derivatization and detection of lipid *N*-chloramines (Figure 1A).

**Figure 1.**
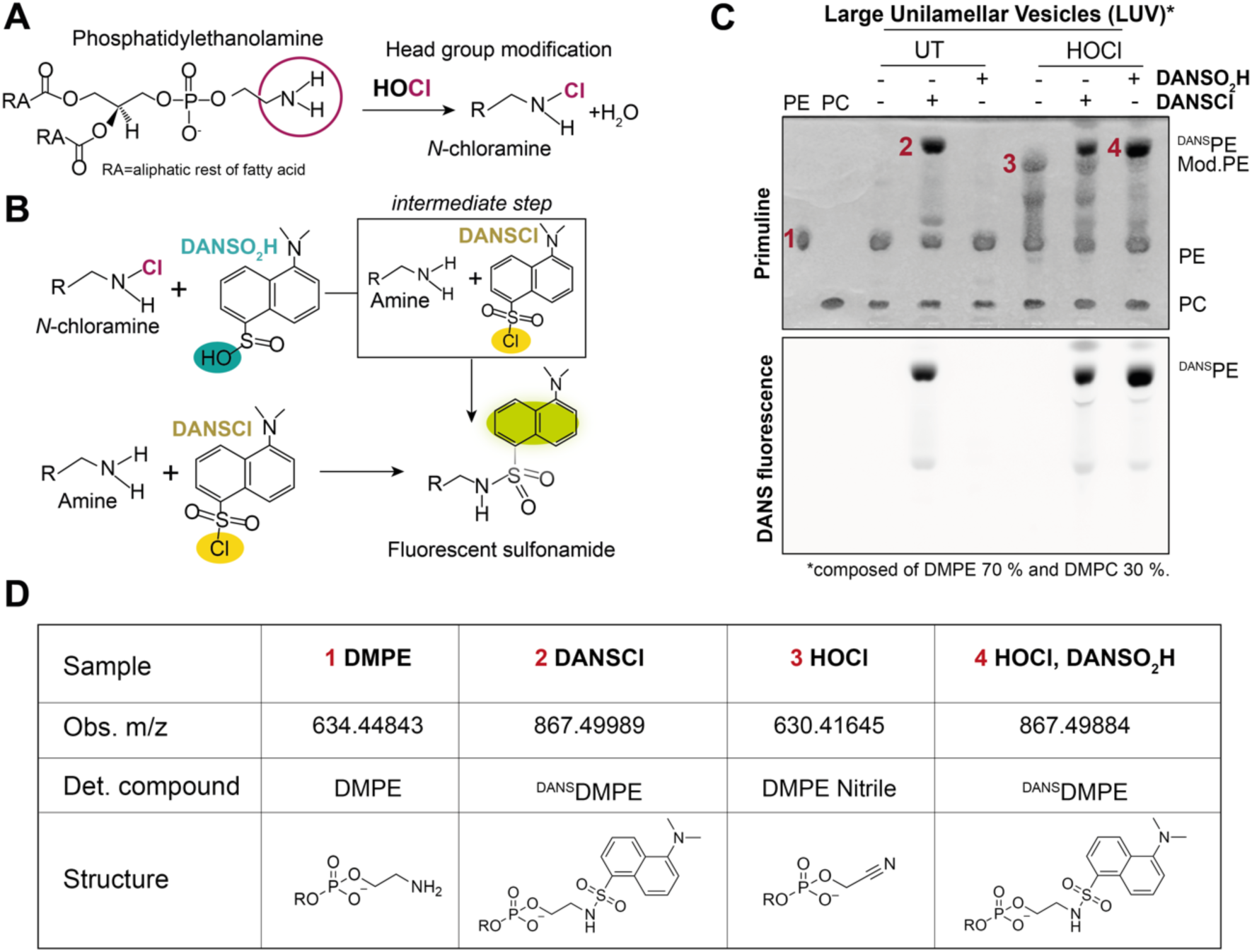
Detection of phosphatidylethanolamine *N*-chlorination with DANSO_2_H in vesicles. **(A)** HOCl-derived *N*-chlorination of the PE head group, resulting in monochloramines. **(B)** Specific derivatization of free amine groups with DANSCl (bottom) and *N*-chloramines with DANSO_2_H (top). DANSO_2_H reacts exclusively with *N-*chloramines forming DANSCl and a free amine group in an intermediate step, ultimately leading to a fluorescent sulfonamide. **(C)** DANSO_2_H derivatization of lipid *N*-chloramines in HOCl-exposed model membranes. Vesicles composed of 70 % PE and 30 % PC (3 mM total lipid) were treated without (UT) or with equimolar HOCl relative to PE (2.1 mM). Excess oxidant was removed prior to derivatization with DANSCl or DANSO_2_H. Lipids were extracted and analyzed by TLC. Chromatograms shown were imaged under UV light before (DANS fluorescence) and after staining with primuline (total lipids). **(D)** LC-MS/MS analysis of head group modifications. For MS/MS analysis, lipids were isolated from the TLC plate in (C) and subjected to LC-MS/MS. MS and MS fragmentation spectra are shown in Figure 2. DMPC, 1,2-dimyristoyl-*sn*-glycero-3-phosphocholine; DMPE, 1,2-dimyristoyl-*sn*-glycero-3-phosphoethanolamine in methanol/chloroform 1/9; LUV, Large unilamellar vesicle; RA, Fatty acid; RO, Phosphoglycerol; R, biomolecule.

To test our hypothesis, we used large unilamellar vesicles (LUV) with a defined lipid composition as model membranes consisting of 70 % PE to mimic the PE content of *E. coli* membranes and 30 % PC (phosphatidylcholine) to support vesicle formation. All lipids used for vesicle formation (PE and PC) contained saturated fatty acids to exclude side chain oxidation by HOCl. The vesicles (3 mM total lipids) were then exposed to a 1:1 molar ratio of HOCl (2.1 mM) relative to the PE content. Excess oxidant was removed by size exclusion and the lipids were isolated by organic extraction. TLC was performed to analyze for the presence of fluorescent sulfonamides formed by the reaction of DANSCl with amines or DANSO2H with *N*-chloramines (Figure 1B). As hypothesized, DANSO_2_H did indeed react with modified PE in HOCl-treated vesicles resulting in the accumulation of fluorescent sulfonamides. However, some head group amines could still be derivatized with DANSCl, indicating that unmodified PE remained even after HOCl treatment. When the vesicles were exposed to HOCl, additional lipid species with different mobility could be detected by TLC, even in the absence of DANSO_2_H. Since these species were not detected after DANSO_2_H derivatization, they may represent *N*-chlorinated PE or degradation products. We used untreated vesicles labelled with DANSCl or DANSO_2_H as specificity control. Only the reaction of DANSCl with these vesicles yielded fluorescent sulfonamides, whereas no reaction was observed with DANSO_2_H, confirming that DANSO_2_H does not react with the unmodified PE head group amine (Figure 1B).

Individual lipid spots (marked with 1-4) in Figure 1B) were re-isolated from the silica gel using organic extraction and subjected to LC-MS/MS detection. The lipid species in spot 2 and spot 4 showed dansyl fluorescence and accumulated after derivatization of untreated vesicles with DANSCl (spot 2) or after HOCl treatment and DANSO_2_H derivatization (spot 4). In both spots (2 and 4), we identified the dansylated PE headgroup using the fragmentation spectra (Figure 1C, Figure 2) indicating the formation of lipid *N*-chloramines upon exposure to HOCl. In spot 3), the lipid species were exclusively found after exposure of the vesicles to HOCl. We first hypothesized that these additional lipid species are chlorinated PE (Figure 1B). However, MS/MS detection failed to detect a chlorine-specific mass shift. Previous studies have shown that PE-*N*-chloramines can decompose to PE nitrile (Kawai et al., 2006) and, indeed, MS/MS fragmentation revealed the presence of PE nitrile, indirectly indicating PE *N*-chlorination. In line with this, the lipid species in spot 3 disappeared when the HOCl-treated vesicles were labelled with DANSO_2_H suggesting that upon HOCl treatment of the vesicles PE *N*-chloramines are formed that can be derivatized with DANSO_2_H.

**Figure 2.**
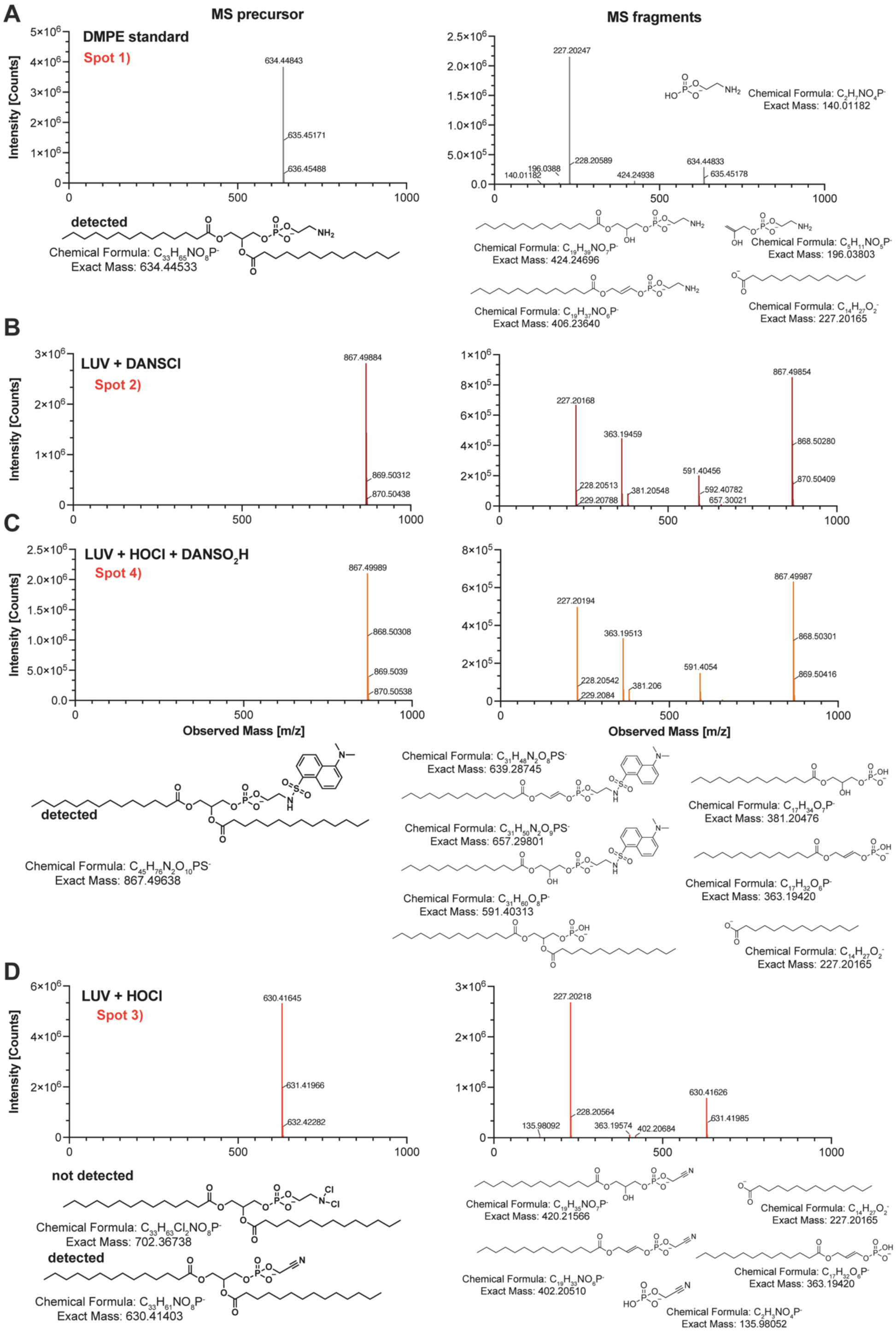
MS/MS precursor and fragmentation spectra and respective structures. DANSO_2_H derivatization of lipid *N*-chloramines in HOCl-exposed model membranes. LUVs composed of 70 mol% DMPE and 30 mol% DMPC were treated with 2.1 mM HOCl and excess oxidant was removed by chromatography prior to derivatization with DANSCl or DANSO_2_H. Lipids were extracted and analyzed by TLC. The marked lipids were isolated by organic extraction from spots 1-4 (refer to Figure 1; spot 1, DMPE standard **(A)**; spot 2, DANSCl-labelled vesicles without HOCl **(B)**; spot 3, vesicles with HOCl **(D)**; and spot 4, vesicles with HOCl and DANSO_2_H **(C)**). Isolated lipids were subjected to LC-MS/MS. MS1 (left, precursor) and MS2 (right, fragments) spectra of the lipid species isolated from the silica plate are shown with the chemical formulas and structures of the identified headgroup. DMPC, 1,2-dimyristoyl-*sn*-glycero 3-phosphocholine; DMPE, 1,2-dimyristoyl-*sn*-glycero 3-phosphoethanolamine; LUV, Large unilamellar vesicles.

In summary, we observed accumulation of dansylated PE after derivatization of HOCl-treated model membranes with DANSO_2_H using TLC and LC-MS. Simultaneously, the band we identified as PE nitrile completely disappeared after DANSO_2_H derivatization, indicating that DANSO_2_H can be used to scavenge highly reactive PE *N-*chloramines.

### Phosphatidylcholine does not react with HOCl

PC is an important membrane lipid for eukaryotic cells, and is also found in some bacteria that interact with eukaryotic hosts. The head group of PC is a choline residue that is synthesized in the cell by methylation of the amino head group of PE to a quaternary ammonium cation (Geiger et al., 2013). The fully methylated head group of PC should not be susceptible to *N*-chlorination (Figure 3A). To test this, we generated PC vesicles (3 mM total lipid) and exposed them to 2.1 mM HOCl. After removal of the oxidant, derivatization with DANSO_2_H and DANSCl was performed as described above. Lipids were isolated and analyzed via TLC, showing that neither DANSCl, nor DANSO_2_H treatment resulted in the formation of dansylated lipids (Figure 3B). In contrast to PE vesicles, exposure of PC vesicles to HOCl did not result in accumulation of additional lipid species or dansylated PE, demonstrating that HOCl does indeed not react with the head group of PC.

**Figure 3.**
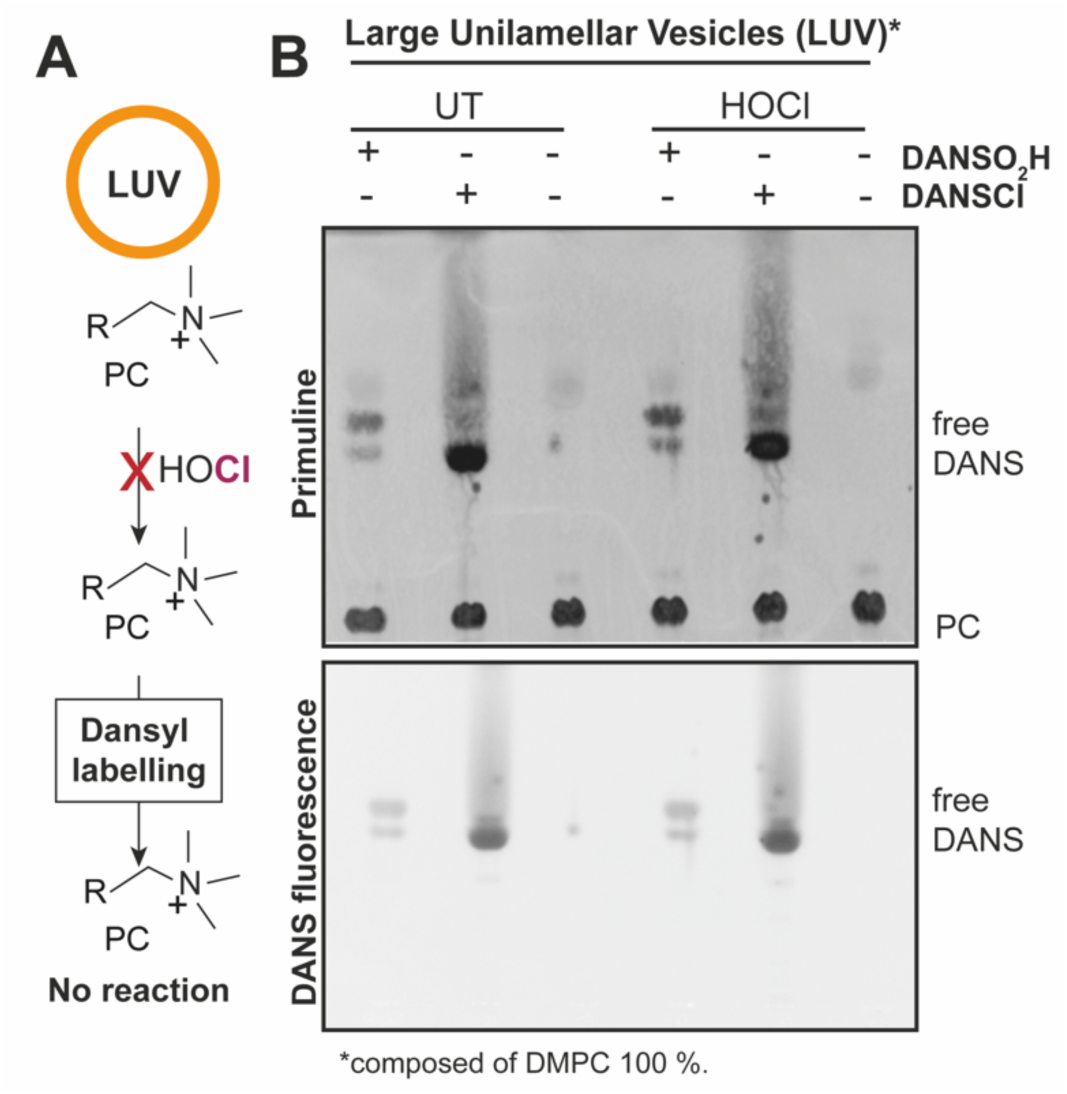
Phosphatidylcholine with fully saturated fatty acids does not react with HOCl or DANSCl/SO_2_H. **(A)** Schematic representation of the workflow. **(B)** Dansyl derivatization of vesicles composed of PC. Vesicles (3 mM) were exposed to HOCl (2.1 mM) or ddH_2_O (UT) and after oxidant removal, derivatization with DANSCl or DANSO_2_H was performed. Lipids were extracted and analyzed by TLC. Chromatograms shown were imaged under UV light before (DANS fluorescence) and after staining with primuline (total lipids). LUV, Large unilamellar vesicles.

### HOCl-driven N-chlorination of the membrane lipid PE is significantly more efficient in E. coli membranes compared to THP1-monocytes

During infection neutrophils can form NET (Neutrophil extracellular traps) structures in which HOCl-producing MPO (myeloperoxidase) is localized outside of the cell. It has been estimated that locally HOCl levels in the millimolar range (1-50 mM) can occur at sites of inflammation (Summers et al., 2008; Weiss, 1989). Therefore, we hypothesize that not only phagocytized bacteria are targeted by the oxidant during infection, but also free bacteria or even host immune cells themselves, such as macrophages.

In order to detect lipid *N*-chloramines in free living cells, we used *E. coli* as model for bacteria and THP1-monocytes as model for mammalian cells. We estimated the theoretical PE amount in *E. coli* (320 µM in a OD_600_ of 6 solution) and THP1 (500 µM in a 10^6^ cells/mL solution) cell solutions (see Materials and Methods). We then exposed the cells to an equimolar amount of HOCl or to a 10-fold molar excess relative to theoretical PE.

To confirm that these HOCl concentrations have an *in vivo* effect, we used *E. coli* expressing the redox sensor roGFP2 in the cytoplasm (Figure 4A). This redox sensor is a GFP variant, in which two cysteines (positions 147 and 204) have been introduced that form a disulfide bond upon oxidation resulting in changes in the excitation spectrum with constant emission at 510 nm. The excitation maximum at 488 nm is reversibly decreased with simultaneous increase of the 405 nm maximum. Therefore, this probe allows a ratiometric (I_405 nm_/I_488_ nm) and concentration independent determination of its dithiol disulfide state (Dooley et al., 2004; Lukyanov and Belousov, 2014; Meyer and Dick, 2010; Müller et al., 2017). Both HOCl concentrations fully oxidized the cytosolic probe within seconds, confirming that HOCl penetrates both the inner and outer membranes of *E. coli* under the conditions used in our *N*-chlorination approach (Figure 4B).

**Figure 4.**
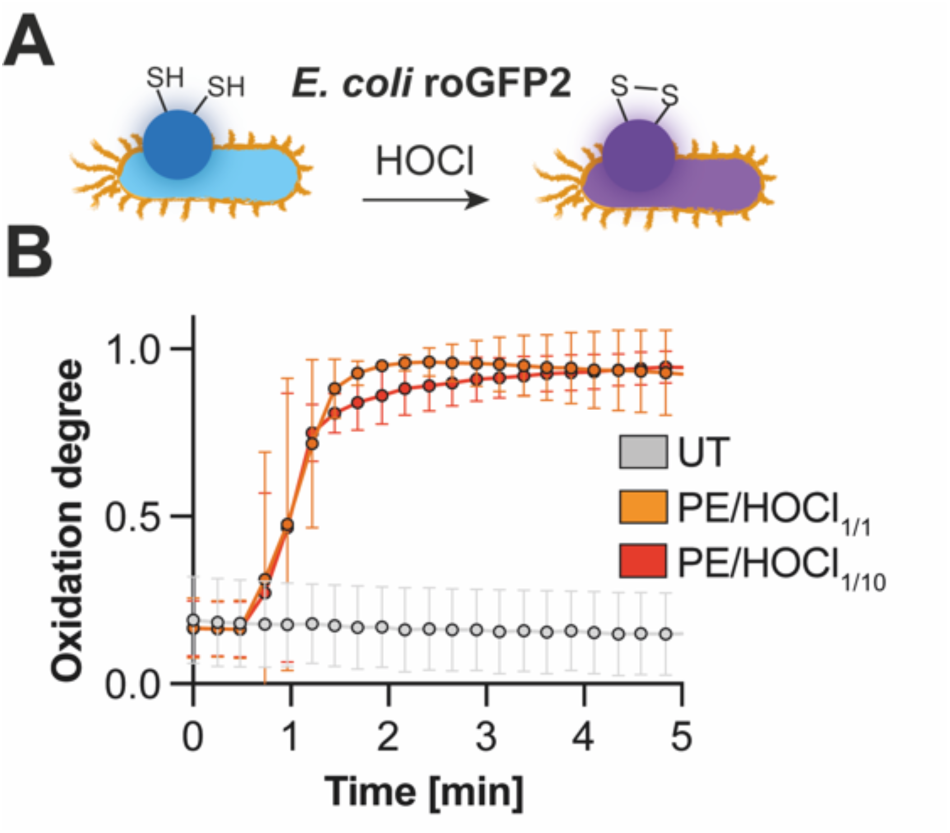
HOCl-driven oxidation of roGFP2 expressed in the cytosol of *E. coli*. **(A)** Schematic representation of the workflow. **(B)** Oxidation of roGFP2 upon exposure to HOCl. roGFP2 expressing *E. coli* cells were treated with an equimolar (1:1) or a 10-fold molar excess of HOCl in relation to theoretical PE content. Untreated cells served as control (UT). Fluorescence was recorded with 510 nm emission and 350 nm to 500 nm excitation. The oxidation degree (OxD) was calculated from the ratio of the fluorescence intensities measured at 405 nm and 488 nm. AT-2 oxidized and DTT-reduced cells served as controls for calculation of OxD. Error bars represent the standard deviation and values (°) are the mean of *n*=3 individual replicates.

Next, we exposed both cell types to an equimolar amount of HOCl or to a 10-fold molar excess, labelled the cells with our dansyl probes and analyzed lipid *N*-chlorination (Figure 5A). As expected, HOCl treatment with 10 x HOCl (millimolar) resulted in lipid *N*-chloramine formation, in both, *E. coli* and THP1-derived monocytes (Figure 5B). Even 1 x HOCl (micromolar) led to lipid *N*-chloramine formation in both cell types, albeit at significantly lower levels compared to higher concentrations. In contrast to *E. coli*, dansylated lipids were barely detectable in THP1 cells after exposure to equimolar HOCl. Compared to *E. coli*, *N*-chlorination seems to be less efficient in monocytes at both HOCl concentrations.

**Figure 5.**
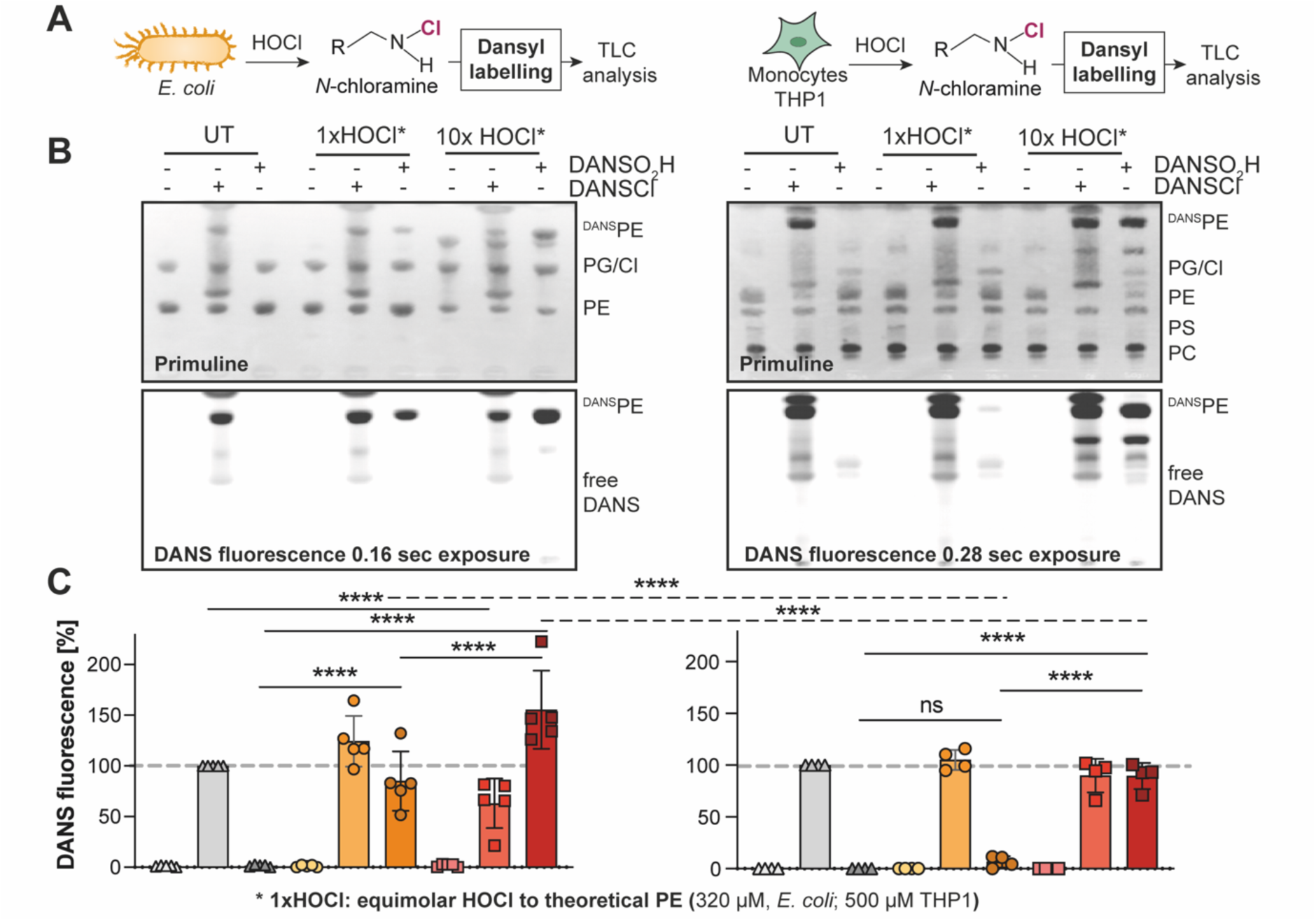
Exposure of *E. coli* and monocytic THP1 cells to HOCl results in formation of PE *N*-chloramines *in vivo*. **(A)** Schematic representation of the workflow. **(B)** Labelling of untreated (UT) or HOCl-treated *E. coli* or THP1 cells with dansyl probes. HOCl treatment for 10 min involved an equimolar (*E. coli*: 320 µM/THP1: 500 µM) or a 10-fold molar excess (*E. coli*: 3.2 mM/THP1: 5 mM) concentrations. After removing excess of HOCl, a two-fold molar excess of dansyl probes relative to PE (*E. coli*: 640 µM/THP1: 1 mM) was added for labelling of free amine groups (DANSCl) or *N*-chloramines (DANSO_2_H). Lipids were then isolated and analyzed by TLC. Chromatograms shown were imaged under UV light before (DANS fluorescence) and after staining with primuline (total lipids). **(C)** Band intensities of dansylated PE were quantified using ImageJ. Dansylated PE of untreated cells incubated with DANSCl were set 100 %. Each value (°) represents one independent experiment and error bars represent the standard deviation of the means of at least *n=4* replicates. *p* values were calculated using one-way ANOVA; \**p*<0.01; \*\*\*\**p*<0.0001.

We used DANSCl derivatization as control for residual PE after HOCl treatment. For both, *E. coli* and THP1, no significant differences in dansylated PE were observed in untreated or 1 x HOCl-treated cells. When 10 x HOCl was used, the total amount of detected free amine groups decreased significantly in *E. coli*, but not in THP1 cells. Untreated cells were labelled with DANSO_2_H to exclude unspecific reactions *in vivo* of our probe with membrane lipids, suggesting that our probe is specific for lipid *N*-chloramines (Figure 5B).

The fact that we were able to observe lipid *N*-chlorination at physiological HOCl concentrations *in vivo* suggests that this modification could play a role during severe inflammation.

### The redox active roGFP2 protein is oxidized by PE N-chloramines in vitro

Previously, model *N*-chloramines such as taurine or lysine *N*-chloramines have been shown to oxidize various thiol-containing plasma proteins such as human serum albumin (HSA). HOCl-modified proteins are associated with several diseases such as atherosclerosis and are involved in the killing of invading pathogens (Summers et al., 2008).

Therefore, we investigated the ability of lipid *N*-chloramines to oxidize thiol-containing biomolecules. For that, we used the redox active model protein roGFP2 (Figure 6A). We reduced purified roGFP2 with DTT, removed the reductant and measured the probés oxidation degree after addition of chlorinated vesicles, prepared as described above.

**Figure 6.**
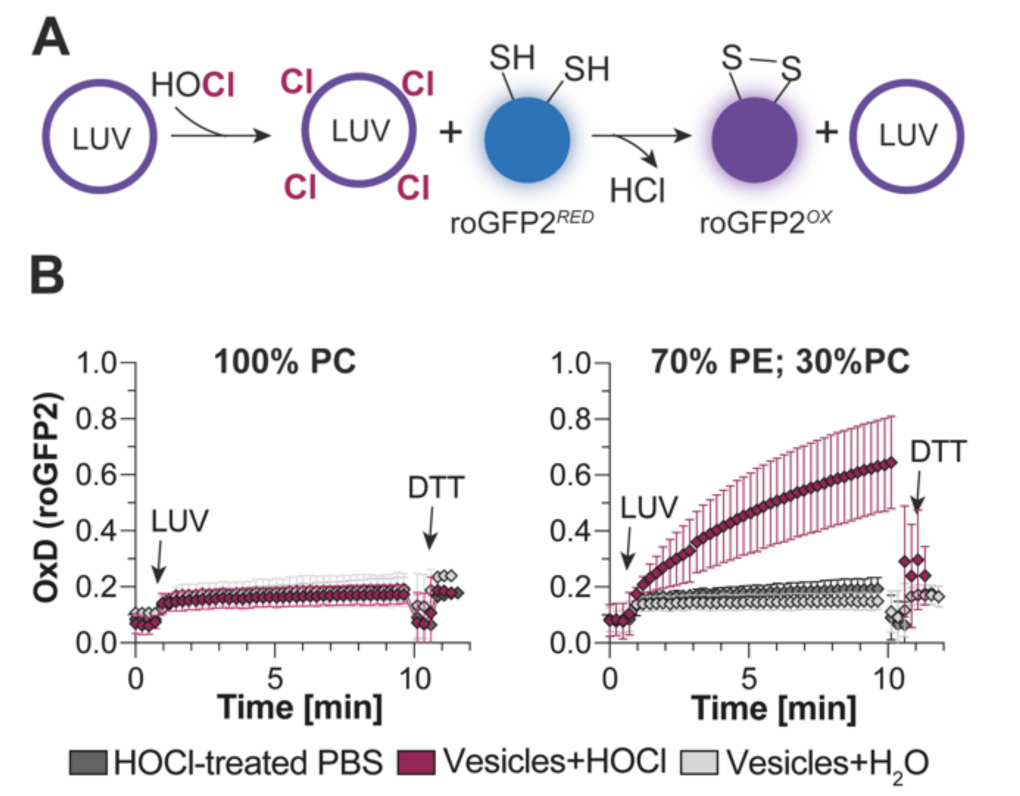
Chlorinated PE vesicles oxidize purified, reduced roGFP2 *in vitro*. **(A)** Schematic workflow. **(B)** roGFP2 oxidation in presence of chlorinated vesicles. Sensor oxidation was tracked by interval scanning of the excitation spectrum (350-500 nm) at 510 nm emission. After 1 min baseline measurement, a 100-fold molar excess of HOCl-treated PC vesicles or PE vesicles (28.6 µM lipids, 20 µM PE) relative to roGFP2 (0.2 µM) was added. Fluorescence was recorded for 5 min. DTT (20 mM) was added to fully reduce the sensor. HOCl-treated PBS, passed over a NAP-5 column, served as control for residual HOCl, while water-treated vesicles were the negative control. The oxidation degree (OxD) of the sensor was calculated from the ratios of the intensities at 405 nm/488 nm. AT-2-oxidized and DTT-reduced roGFP2 served as controls for calculation of OxD. Values are the mean of *n=4* independent replicates and error bars represent the standard deviation. LUV, Large unilamellar vesicles.

Chlorinated vesicles containing 70 % PE were able to oxidized our roGFP2 redox probe within minutes (Figure 6B). To account for residual HOCl after oxidant removal, we used HOCl-treated PBS buffer that was passed through a NAP-5 column. The chromatography column used was capable of completely removing excess HOCl, as evidenced by the inability of HOCl-treated PBS to oxidize roGFP2. To exclude HOCl transport by the vesicles, we used vesicles composed of 100% PC. Since PC itself is not *N*-chlorinated by HOCl (Figure 3), these vesicles should not exhibit oxidative activity, and indeed roGFP2 was not oxidized in the presence of HOCl-treated PC vesicles (Figure 6B).

Taken together, our data suggests that lipid *N*-chloramines are part of the deleterious cocktail of HOCl-modified biomolecules generated during infection and inflammatory diseases and could mediate the killing of invading pathogens.

### N-chlorination of phosphatidylethanolamine can be reversed by glutathione

Cells have evolved various reduction systems to cope with oxidation and keep their cytosolic proteins reduced. One of the most abundant cellular reductants is the low molecular weight thiol glutathione (GSH) which is present at levels of up to 5-10 mM in a wide range of cells, including bacteria and mammalian cells (Fahey et al., 1978; Meister, 1988; Vašková et al., 2023). Cytosolic GSH is actively kept in its reduced state by the NADPH-dependent glutathione reductase. Glutathione is also present in the periplasm of Gram-negative bacteria, however in a more oxidized state and at lower concentrations. Upon oxidation, GSH dimerizes by formation of a disulfide, also referred to as GSSG (Carmel-Harel and Storz, 2000; Knoke et al., 2023; Masip et al., 2006; Smirnova et al., 2012). Previous studies have shown that GSH not only reduces protein thiols but also reverses *N*-chlorination of amino acids (Carroll et al., 2017; Koga et al., 2024; Müller et al., 2014; Peskin and Winterbourn, 2003, 2001).

To analyze if GSH can repair lipid *N*-chloramines, we again generated PE vesicles (3 mM total lipid). The vesicles were exposed to an equimolar amount of HOCl (2.1 mM) in relation to the PE concentration and excess oxidant was removed. Chlorinated vesicles were reduced with a 10-fold molar excess of DTT or GSH relative to PE, followed by subsequent dansyl derivatization (Figure 7A). Untreated and non-reduced vesicles served as controls for *N*-chloramine formation and DTT-treated vesicles as controls for the reversibility of PE *N*-chloramine formation by thiol-based reductants. After dansyl derivatization, lipids were isolated, dansylated PE and total lipids were visualized using TLC and spot density of dansylated PE was quantified using ImageJ.

**Figure 7.**
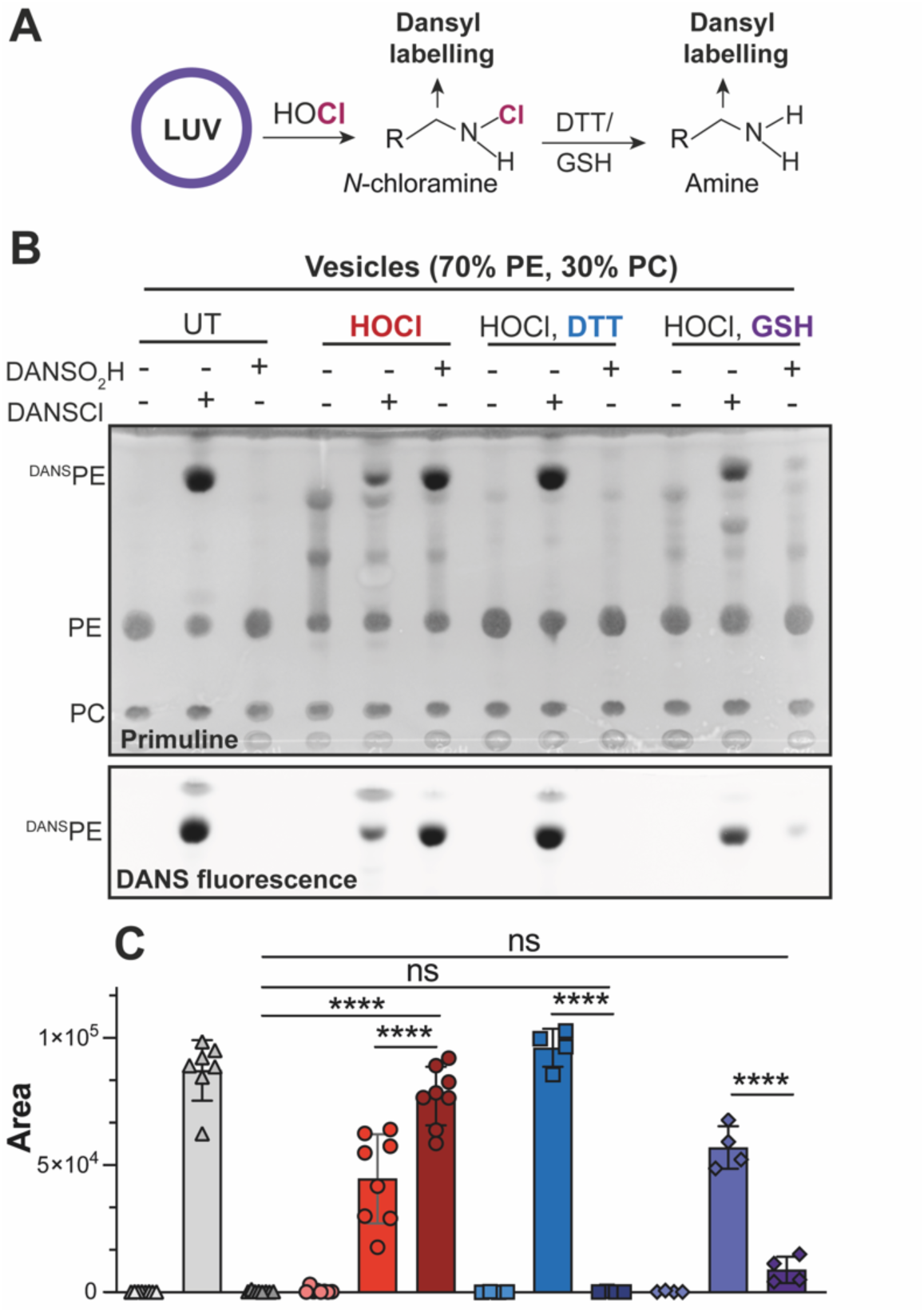
Glutathione reverses *N*-chlorination of PE in model membranes. **(A)** Schematic representation of the workflow. **(B)** Thin layer chromatography of isolated lipids. PE vesicles (1500 nmol total lipid) were treated with equimolar HOCl in relation to the PE concentration (1050 nmol). Excess HOCl was then removed by size exclusion chromatography. 517 nmol lipids were incubated with a 10-fold excess of GSH or DTT (5.17 µmol). After that, reductants were removed. Dansyl labelling was performed with all indicated samples with 100 nmol total lipid each. For that, a two-fold molar excess of dansyl probes (140 nmol) in relation to PE (70 nmol) was used. Untreated vesicles (UT) served as reference for free amines and non-labelled vesicles as negative control. After labelling, lipids were isolated. Chromatogram shown was imaged under UV light before (DANS fluorescence) and after staining with primuline (total lipids). **(C)** Quantification of dansylated PE. For quantification, band intensities of dansylated PE were quantified from the dansyl fluorescence image using ImageJ. Each value (°) represents one independent experiment and error bars represent the standard deviation of the means of at least *n=4* replicates. *p* values were calculated using one-way ANOVA; \**p*<0.01; \*\*\*\**p*<0.0001. LUV, large unilamellar vesicle.

We again observed *N*-chloramine formation when vesicles were exposed to HOCl with a concomitant decrease in labelled PE amines. *N*-chlorination of PE-containing vesicles was reversed by glutathione. However, very small amounts of *N*-chlorinated PE remained (Figure 7B). In contrast, treatment of chlorinated vesicles with DTT completely reversed *N-*chlorination of PE.

## Discussion

Professional phagocytotic cells, such as neutrophils, produce the enzyme myeloperoxidase (MPO), which catalyzes the formation of the highly toxic oxidant hypochlorous acid (HOCl), important for killing of engulfed pathogens (Sultana et al., 2020; Ulfig and Leichert, 2021). In addition to oxidation of protein thiols (Xie et al., 2019), HOCl attacks amine groups in various target molecules such as lysine residues in proteins, carbohydrates, nucleic acids and lipids, resulting in *N*-chlorination (Panasenko et al., 2013; Schröter and Schiller, 2016; Varatnitskaya et al., 2022). Previous studies have shown that HOCl induces *N*-chlorination in lipids containing a head group amine, such as phosphatidylethanolamine (PE) (Carr et al., 1998; Jaskolla et al., 2009; Richter et al., 2008), but its occurrence *in vivo* has not yet been demonstrated. Since it has been estimated that HOCl levels at the site of infection can rise to 50 mM (Summers et al., 2008), we hypothesized, that lipid *N*-chloramines are formed at physiological HOCl concentrations.

Here, we used the dansyl derivative DANSO_2_H in combination with lipid analysis to detect lipid *N*-chloramines at physiologically relevant HOCl concentrations in both *E. coli* and monocytic cells. HOCl induced *N*-chlorination of THP1 membranes was significantly less effective compared to *E. coli* membranes especially at lower concentrations. This observed difference could be explained by the subcellular distribution of lipids in eukaryotic cell compartments. While PC is the most prominent lipid in the outer leaflet of the eukaryotic plasma membrane, PE is located in the inner leaflet with a fraction of up to 20% and is even more enriched in mitochondrial membranes, where it constitutes up to 40 % of the phospholipids (Ali and Szabó, 2023; Yang et al., 2018). Thus, PE in bacterial membranes could be more susceptible to modification by HOCl in comparison to eukaryotic PE due to its exposure on the inner leaflet of the bacterial outer and the plasma membrane. When formed *in vivo* during phagocytosis, PE might exhibit a protective role by scavenging HOCl before it can enter the cytosol of the pathogen. Alternatively, PE *N*-chloramines might be important for killing of the engulfed pathogen by potentiating the oxidative effect of HOCl.

Indeed, it has been suggested that HOCl-derived modifications are important for killing of invading pathogens and initiating of various diseases such as atherosclerosis, sepsis and rheumatoid arthritis. Small and protein *N*-chloramines are thought to contribute to these deleterious effects by oxidizing thiol-containing molecules and initiating lipid peroxidation (Gaut et al., 2001; Kawai et al., 2006; Summers et al., 2008; Thomas et al., 1986). In addition, chlorinated HSA and other serum components have been shown to activate immune cells and stimulate the immune system (Ulfig et al., 2021, 2019). Based on *in vitro* studies, it has been proposed that *N*-chlorinated PE can induce lipid peroxidation in EYPC (egg-yolk phosphatidylcholine) through dichloramine and *N*-centered radical intermediates, which may enhances cytotoxicity important during inflammation (Kawai et al., 2006). Consistent with the abovementioned studies on protein and model *N*-chloramines, chlorinated vesicles containing PE were able to oxidize the roGFP2 redox-sensing protein. We propose a similar immune modulatory function of *N*-chlorinated lipids as shown for protein *N*-chloramines. Another phospholipid prone to *N*-chlorination is phosphatidylserine (PS), which exclusively resides in the cytoplasmic leaflet of eukaryotic plasma membranes. Its exposure on the cell surface is linked to cell death and inflammation (Flemmig et al., 2009; Flemmig and Arnhold, 2010; Kawai et al., 2006). Concomitantly, we propose a similar function for PS *N*-chloramines in immune modulation as for PE.

Cells have evolved a variety of cellular antioxidant systems to prevent permanent damage of biomolecules by oxidative modifications, such as the thioredoxin or glutathione in combination with the glutaredoxin system (Jena et al., 2023). The tripeptide GSH is present in the millimolar range in the cytosol and actively excreted (Knoke et al., 2023; Pittman et al., 2005; Smirnova et al., 2012). Previous studies using the “chloramine T” model have shown that thiol-containing molecules such as GSH or DTT are oxidized in the presence of *N*-chloramines (Carr et al., 2001; Peskin and Winterbourn, 2003, 2001) suggesting that the cellular GSH pool protects cells from overoxidation by HOCl and its chlorinated products. Indeed, GSH has been shown to protect cells from the toxic effects of monochloramine and taurine *N-*chloramines. During oxidation, GSH forms a dimer called GSSG. Interestingly, the oxidized glutathione dimer also protected cells from damage by HOCl or model chloramines, but the exact mechanism is still under investigation (Chesney et al., 1996). Using chlorinated model membranes, we have shown that GSH detoxifies not only small model chloramines such as taurine chloramine or proteins, but also PE-*N*-chloramines. Possibly, in the periplasm, GSSG/GSH, could protect the cells against HOCl and HOCl-derived chlorine-based oxidants by preventing PE *N*-chloramine induced oxidation of biomolecules and lipid peroxidation. HOCl can cross lipid bilayers in its hydrated form (Kawai et al., 2006) suggesting that lipids in the inner leaflet can also be chlorinated. DTT can cross membranes (Cline et al., 2004), whereas GSH does not (Oestreicher and Morgan, 2019). Our data showing residual PE *N*-chloramines supports this notion, suggesting that residual PE *N*-chloramines are located at the inner leaflet of the vesicles. Another possible reason for the observed residual *N-*chlorination in GSH-treated vesicles could be the difference in the redox potential of the two compounds.

This study, by enabling the stabilization of lipid *N*-chloramines and their detection in living cells, has established a foundation for future investigations targeting the *in vivo* detection of *N*-chloramines during phagocytosis. Lipid *N*-chloramines derivatized with DANSO_2_H hold promise as a novel marker for (chronic) inflammation. Chlorinated model membranes will be instrumental in dissecting their immune modulatory potential and the cytotoxicity of lipid *N*-chloramines, thus advancing our comprehension of their (patho-)physiological roles.

## Acknowledgements

LRK thanks Lars I. Leichert for the generous support of this work and suggestions on this manuscript.

## Author contributions

LRK designed the study, and planned and performed most of the experiments. SAH and TGP helped LRK in planning and SAH in performing model membrane experiments. SH performed mass spectrometry of isolated lipids and NL purified roGFP2. LRK carried out the data interpretation and wrote the manuscript. JEB provided tools. All authors consulted on the manuscript and approved the final version.

## Data availability statement

The data supporting the findings of this study is presented within the article and its supplementary materials.

